# Co-enrichment of cancer-associated bacterial taxa is correlated with immune cell infiltrates in esophageal tumor tissue

**DOI:** 10.1101/2023.05.29.542596

**Authors:** KL Greathouse, JK Stone, AJ Vargas, A Choudhury, N Padgett, JR White, A Jung, CC Harris

**Affiliations:** Department of Biology, Baylor University, Waco, TX; Nutrition Division, Human Sciences and Design, Baylor University, Waco, TX; Center for Cancer Research, National Cancer Institute, Bethesda, MD; Eunice Kennedy Shriver National Institute of Child Health and Human Development, National Institutes of Health, Bethesda, MD; Harvard T.H. Chan School of Public Health, Harvard University, Boston, MA; Resphera Biosciences, LLC, Baltimore, MD

## Abstract

Esophageal carcinoma (ESCA) is a leading cause of cancer-related death worldwide, and Barrett’s esophagus (BE) is a strong risk factor along with smoking. Smoking is well-known to induce microbial dysbiosis and we asked if BE and esophageal microbiomes had shared microbial alterations that could provide novel biomarkers. We extracted DNA from BE tissues (n=5) and tumors of 158 patients in the NCI-MD case control study and sequenced the 16S rRNA gene (V3-4), with TCGA ESCA RNAseq (n = 173) and WGS (n = 139) non-human reads used as validation. We identified four taxa, *Campylobacter, Prevotella, Streptococcus*, and *Fusobacterium* as highly enriched in esophageal cancer across all cohorts. Using SparCC, we discovered that *Fusobacterium* and *Prevotella* were also co-enriched across all cohorts. We then analyzed immune cell infiltration to determine if these dysbiotic taxa were associated with immune signatures. Using xCell to obtain predicted immune infiltrates, we identified a depletion of megakaryocyte-erythroid progenitor (MEP) cells in tumors with presence of any of the four taxa, along with enrichment of platelets in tumors with *Campylobactor* or *Fusobacterium*. Taken together, our results suggest that intratumoral presence of these co-occurring bacterial genera may confer tumor promoting immune alternations that allow disease progression in esophageal cancer.

## Introduction

Barrett’s esophagus (BE) is a proinflammatory condition which is a well-established risk factor for development of esophageal adenocarcinoma (EAC), the major form of esophageal cancer (ESCA). Incidence rates of ESCA have increased in recent decades in Europe and North America with rates predicted to rise [1-3]. The histopathological changes resulting in development of ESCA can result from a range of environmental and genetic factors, including obesity, tobacco smoking, and *TP53* mutations [4-7]. These risk factors are also known to play a role in modulating the gastrointestinal microbiome [8].

Several studies have demonstrated community and taxonomic alterations of the esophageal microbiome in BE or EAC patients [9-11], showing a transition from Gram-positive dominated to a Gram-negative dominated microbiota in BE. *In vivo* studies indicate that microbiome changes occur during the development of BE and EAC that correlate with changes in gene expression in the esophageal epithelium, including multiple microbial sensing pathways (e.g. toll-like receptors) that influence immune signaling and immune cell recruitment patterns [12, 13]. These data suggest that alterations in the microbiome in development of BE may result in chronic inflammation and contribute the to the development of EAC.

## Results

To better understand the microbial and immune system contributions to BE and ESCA development, we comprehensively evaluated the esophageal tissue microbiome and inferred immune cell infiltration in patients with BE or ESCA (with and without a history of BE) in two cohorts, National Cancer Institute-Maryland (NCI-MD) and The Cancer Genome Atlas (TCGA) whole genome sequencing (WGS), and TCGA RNA sequencing (RNA-seq), which results in three datasets. We extracted DNA from patients enrolled in the NCI-MD case control study from the Baltimore, Maryland area (126 non-tumor tissues, 5 BE, 98 tumors; 45 NT-T pairs) for 16S V3-4 sequencing as previously described (Supplemental Methods) [14]. Briefly, sequence reads were filtered for length (>200bp) and max error rate (0.5%), and submitted for high-resolution taxonomic assignment (Resphera Insight) to assess taxa abundance (Supplemental Methods, Date File 1). TCGA ESCA RNA-seq data (11 non-tumor tissue, 162 tumors; 11 NT-T pairs) and whole genome sequencing (WGS) (61 non-tumor tissue, 62 tumors; 61 NT-T pairs) data were downloaded (GDC Data Portal, NCI) as validation cohorts (Figure 1A, Tables S1-S3). Stringent quality control measures were applied on both data sets (Supplemental Methods, Data File 1).

**Fig. 1:**
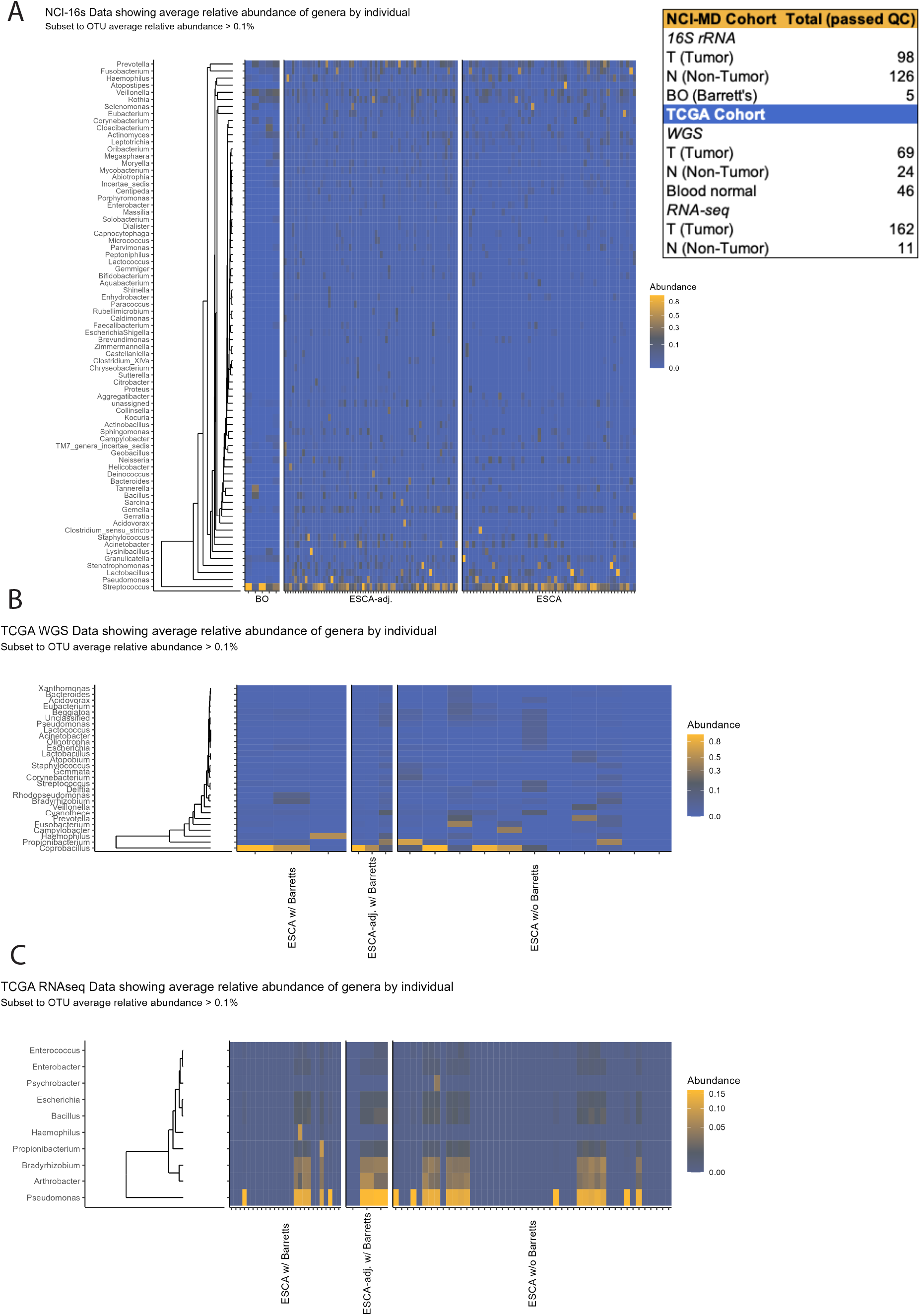
Identification of microbial signatures in Barrett’s esophagus and esophageal cancer. **A** Bacterial abundance within the NCI-MD case control study calculated from 16S V3-4 amplification. EAC adj. w/ Barrett’s indicates non-tumor, adjacent tissue with Barrett’s esophagus as a comorbidity. **(Panel)** Total number of patients used in this study from three cohorts: NCI-MD case control study, TCGA RNA-seq, and TCGA whole genome seq (WGS). **B** Bacterial abundance within TCGA WGS, determined by quantification of non-human aligned reads. **C** Bacterial abundance within TCGA RNA-seq, determined by quantification of non-human aligned reads.

First, we sought to determine if any differences existed in the microbiome between ESCA tumor and non-tumor adjacent tissues with and without BE history. No differences in alpha or beta diversity were seen for the NCI-MD cohort, however alpha diversity decreased significantly in TCGA RNA-seq tumor samples but increased significantly in TCGA WGS tumor samples (Figure S1). Because tobacco smoking is a key risk-factor for developing esophageal cancer [15], we asked if smoking status or other key ESCA risk factors were associated with alpha diversity. Interestingly, none of these factors (gender, histology, race, smoking status, or stage) showed significant difference in taxa abundance across all three cohorts (Figure S2).

Examination of the most abundant taxa in the NCI-MD cohort, independent of tissue type, identified *Streptococcus, Pseudomonas, Prevotella, Veillonella, Lactobacillus, Stenotrophomonas, Fusobacterium*, and *Acinetobacter*. One (*Pseudomonas*) and six (*Streptococcus, Pseudomonas, Prevotella, Veillonella, Lactobacillus*, and *Fusobacterium*) of these taxa were also highly abundant in TCGA RNA-seq and WGS cohorts, respectively (Figure 1). We then performed a statistical concordance analysis (Supplemental Methods), which identified four common taxa across all cohorts: *Campylobacter, Fusobacterium, Prevotella*, and *Streptococcus* as enriched in ESCA (Figure 2). All four taxa were enriched in tumors in at least two of three cohorts (Table S4). Additionally, given that *TP53* is one of the most frequently mutated genes in ESCA [5, 7], we investigated *TP53* mutation status in the TCGA cohort (WGS and RNA-seq) and found no relationship with abundance of these four taxa (Figure S3). To determine if any relationship existed with microbial function, apart from community structure, we investigated the inferred metabolic profile of the ESCA microbiome using PICRUSt, but did not identify any associations with ESCA between tumor vs non-tumor tissue, overall or stratified by the four taxa (data not shown).

**Fig. 2:**
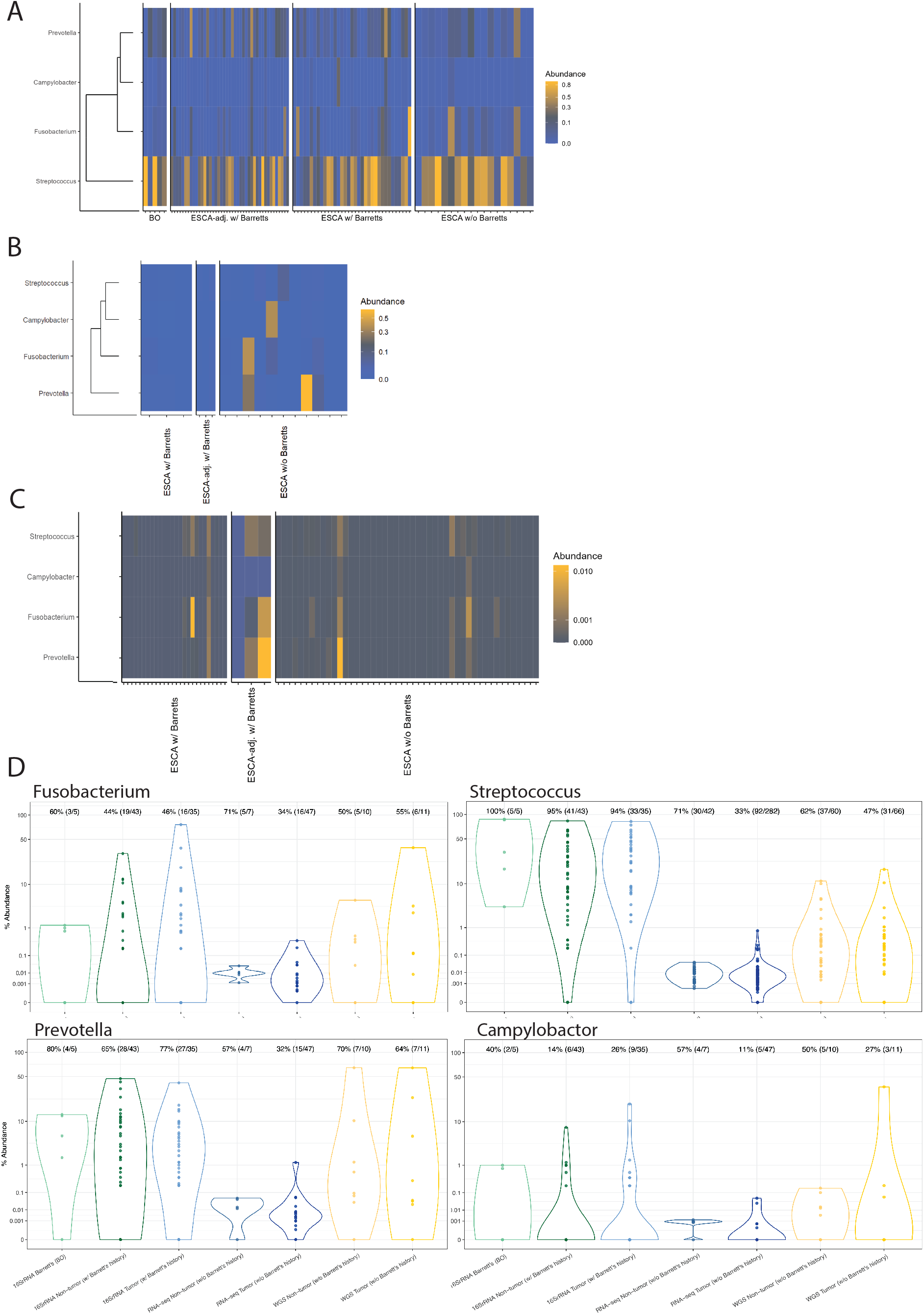
Four taxa are enriched in Barrett’s esophagus and esophageal cancer across cohorts. **A** Abundance of *Campylobacter, Fusobacterium, Prevotella*, and *Streptococcus* within NCI-MD case control study calculated from 16S V3-4 amplification. EAC adj. w/ Barrett’s indicates non-tumor, adjacent tissue with Barrett’s esophagus as a comorbidity. Co-association was determined by statistical concordance analysis (Supplemental Methods). **B** Abundance of the above four taxa within TCGA WGS, determined by quantification of non-human aligned reads. **C** Abundance of the above four taxa within TCGA RNA-seq, determined by quantification of non-human aligned reads. **D** Comparison of the above four taxa in non-tumor adjacent and tumor tissues in each of the three cohorts (NCI-MD, TCGA RNA-seq, and TCGA WGS, left to right). Violin plots indicate relative abundance of each of the four taxa; % is the number of tumors with taxa present (ratio is number of samples with taxa present over number of total samples). *Statistical concordance analysis in Table S4.

We then sought to determine taxa similarity between patients with BE only (BO) as compared to those with ESCA, however due to a paucity of available BO tissue (NCI-MD n = 5, TCGA n = 0), we were unable to make statistically valid comparisons between these groups; though similar trends in enrichment of these four taxa in tumor versus BO were observed between these tissue types (Figure 1 and 2A).

Having observed *Fusobacterium* as one of the most enriched taxa in ESCA tissue, we assessed whether this genus is co-abundant with specific taxa in ESCA. Specifically, *Prevotella*, and *Streptococcus* are often found in oral biofilms alongside *Fusobacterium nucleatum* where the two species rely on *F. nucleatum* binding to salivary protein anchors (e.g. Statherin) and sharing nutrients to grow [16]. Furthermore, *Fusobacterium* and *Prevotella* have been previously described as co-enriched in ESCA [10, 17]; therefore we asked if any of our four enriched taxa (*Campylobacter, Fusobacterium, Prevotella*, and *Streptococcus*) were co-enriched in the same tumors or were associated with other taxa. Using SparCC to compensate for correlation deficiencies inherent to low-level biomass associated with 16S-based studies [18], we calculated co-enrichment for each of the three datasets. Overall, TCGA (RNAseq and WGS) results were highly concordant (Table S5), and showed a consistent co-enrichment of certain taxa, with *Fusobacterium* and *Prevotella* co-enriched across all cohorts (Figure 3A-C, Figure S4). These two taxa were also enriched with *Leptotrichia* and *Veillonella*, consistent with prior reports in colorectal cancer [19, 20]. The NCI-MD cohort beta diversity suggested, however, that *Streptococcus* enriched samples were divergent from those with *Campylobacter, Fusobacterium*, and *Prevotella* (Figure S1E). We also confirmed a negative co-enrichment between *Streptococcus* and *Campylobacter* and *Fusobacterium* in this cohort (Figure 3A).

**Fig. 3:**
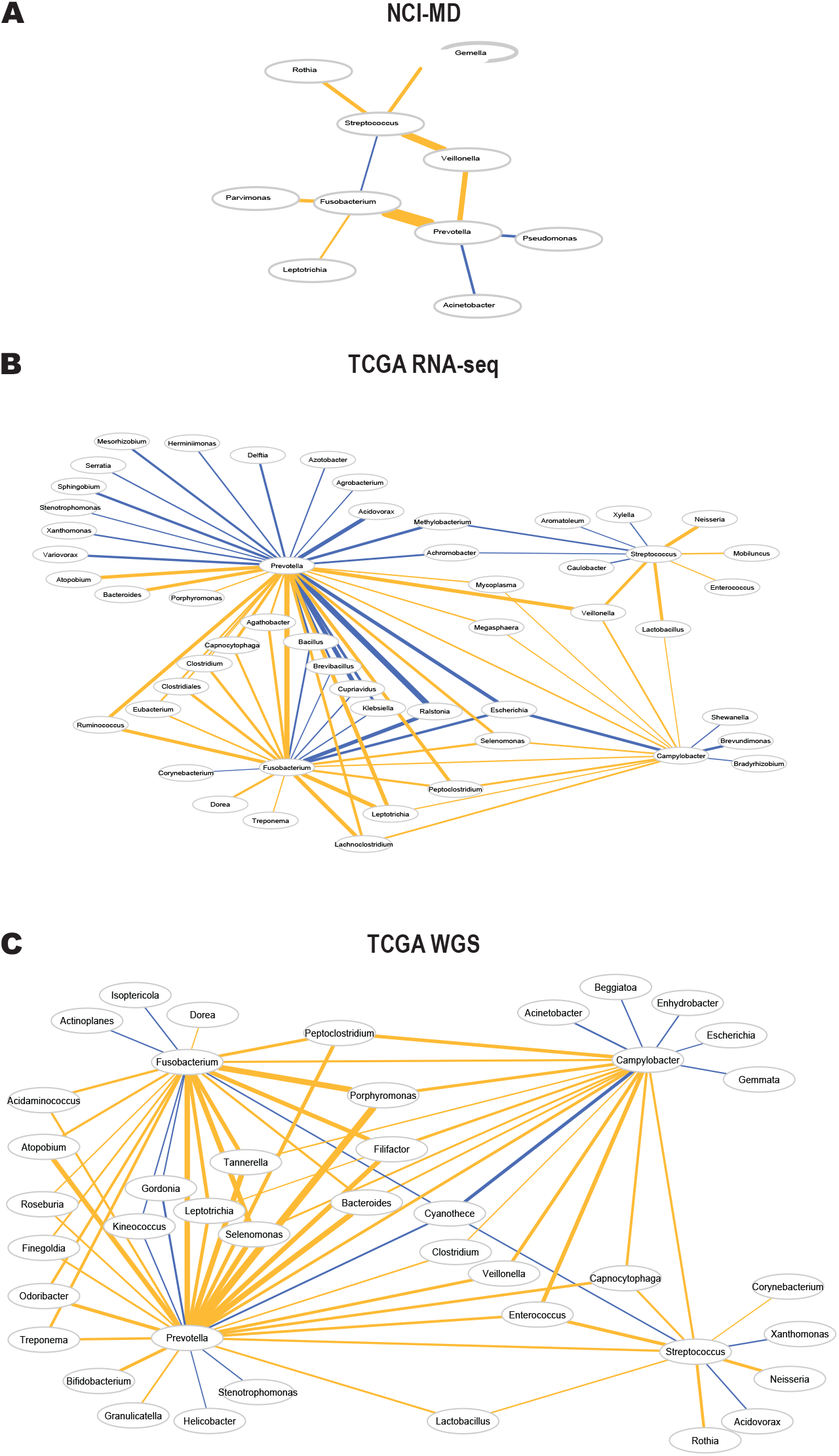
*Fusobacterium* and *Prevotella* are consistently co-associated across cohorts. **A** Taxa co-enrichment networks within the NCI-MD case control study. Taxa abundance was permuted through SparCC with 100 iterations, and correlation coefficients were filtered for X < -0.2 and X > 0.2. Gold edges indicate positive coefficients demonstrating co-enrichment while blue edges indicate negative coefficients demonstrating exclusion. Edge thickness represents normalized coefficient values. **B** Taxa co-enrichment networks within TCGA RNA-seq. **c** Taxa co-enrichment networks within TCGA WGS. Networks for **b** and **c** were filtered for correlation coefficients X < -0.3 and X > 0.3, otherwise networks were constructed as described for **A**.

Regardless of cohort, we found these four taxa were negatively associated with *Acinetobacter, Brevundimonas, Klebsiella, Pseudomonas*, and *Xanthomonas* (Figure 3A-C). These data indicate that co-occurrence of *Fusobacterium* and *Prevotella* are a common feature of ESCA and may be important in ESCA pathology.

Since *Fusobacterium* spp. have demonstrated effects on gene expression changes within tumor epithelial and immune cells, we predicted immune cell infiltration from TCGA RNA-seq data using the deconvolution algorithm xCell [21], and then compared their abundances with our four co-enriched taxa. We identified a depletion of megakaryocyte-erythroid progenitor (MEP) cells in tumors with presence of any of the four bacteria (Figure 4A-B), which was significant in the RNA-seq TCGA dataset (p<0.001) (Figure S5). Furthermore, we found a modest enrichment of platelets in tumors with *Campylobactor* or *Fusobacterium* (p<0.06) (Figure 4A-B, Figure S5). These data suggest that intratumoral presence of these bacterial genera result in loss of MEPs by promoting their terminal differentiation to platelets.

**Fig. 4:**
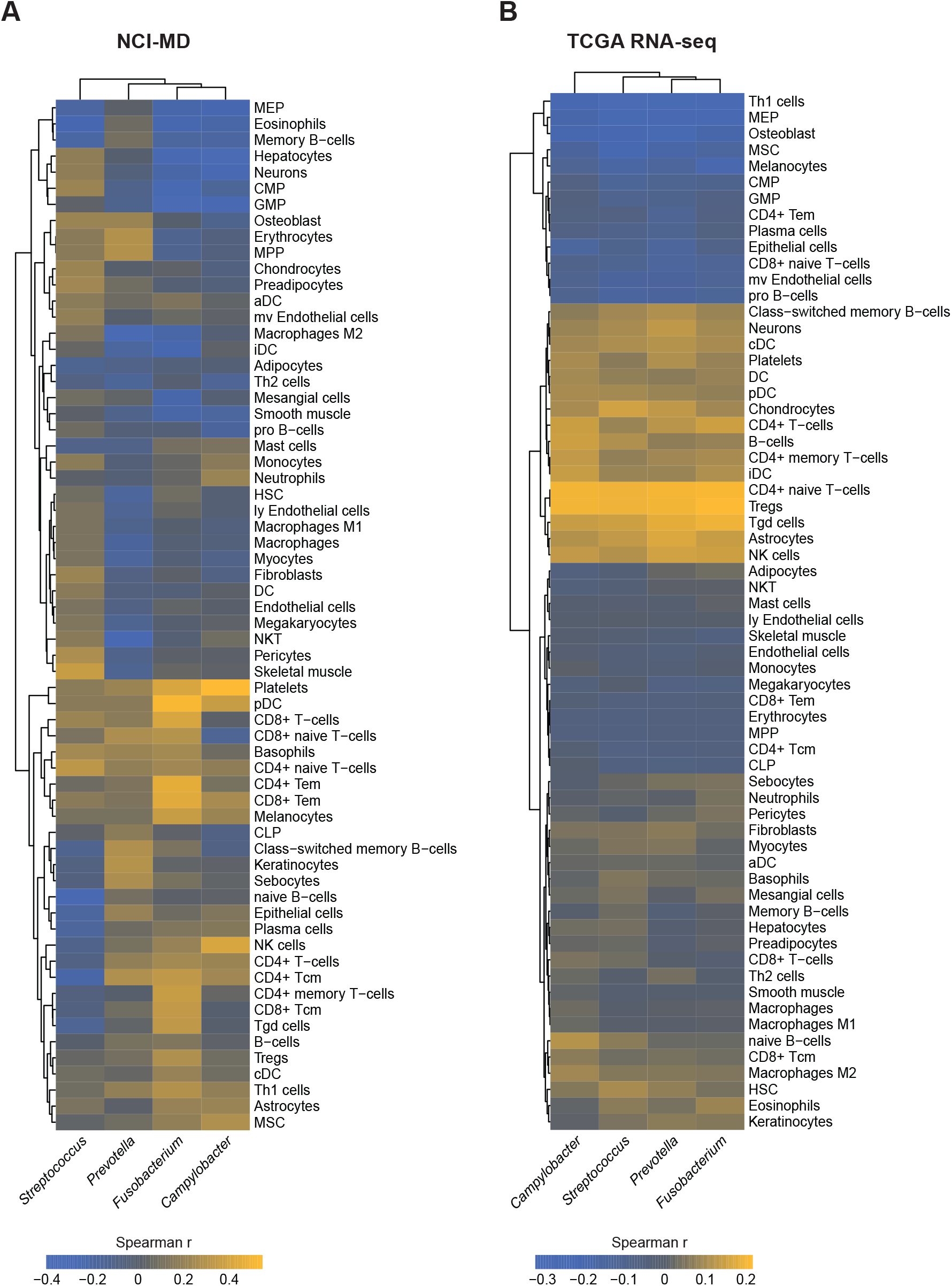
Megakaryocyte–erythroid progenitor cells are depleted in tumors with high carriage of ESCA-enriched taxa. **A** RNA-sequencing was performed on NCI-MD patients (n = 27; non-tumor = 13, BO = 4, tumor = 10) and samples were analyzed for predicted cell infiltration using xCell (citation). Cell infiltrates and taxa abundance were correlated using Spearman’s coefficient. **B** Correlation of xCell predicted cell infiltration in TCGA RNA-seq patients with taxa abundance. *Statistical significance analysis in Fig. S5.

## Discussion

Globally rising rates of ESCA suggest that potentially novel drivers are partially responsible, however identification of these factors remains a significant problem in diagnosis and treatment. In this study we asked if the microbiome, a known driver of various GI malignancies [22], was altered in ESCA in comparison to non-tumor adjacent esophageal tissue. We found enrichment of the taxa *Campylobacter, Fusobacterium, Prevotella*, and *Streptococcus* in tumor tissue. These findings are consistent with other studies examining the ESCA microbiome [9], although our study is the first to report the enrichment of all four taxa in the same cohorts.

Additionally, as these taxa exist within a community, abundance changes in one taxa are associated with changes in others, and we found consistent co-enrichment networks with the above four taxa common across taxa, including associations of *Fusobacterium, Prevotella, Leptotrichia*, and *Veillonella*. Interestingly, these taxa may also invoke terminal differentiation of MEPs into platelets within the tumor microenvironment, possibly due to upregulation of inflammatory signals in MEP cells or bacterial-induced platelet activation [23, 24]. High expression of platelet-derived growth factor A (PDGFA) is a poor prognostic factor in ESCA and platelet expansion or activation may then confer disease progression to metastasis, as occurs in colorectal cancer [25, 26].

These findings suggest that cancer therapeutic strategies targeting one genus or species are likely to fail as other associated taxa may offset the loss of those taxa. Instead, it is likely a multi-targeted strategy, based on presence of intratumoral taxa, for modulating microbial dysbiosis is required to improve treatment and patient outcome. Further research is needed, however, to better understand the mechanisms driving enrichment of these taxa and immune cells in EAC and other cancers.

## METHODS

### University of Maryland (UMD) Esophageal Samples

DNA was extracted from 362 esophageal tissue samples, one plain water control (Mo Bio Laboratories, inc, Carlsbad, CA, USA), one water control (Mo Bio Laboratories, inc, Carlsbad, CA, USA) that was carried through the DNA extraction process, and one mock community (BEI resources, Manassas, VA, USA). Esophageal tissue was collected at the University of Maryland Medical Center (Baltimore, MD, USA) under an IRB-approved collection protocol (OH98CN027/ FWA00005897) where all surgical subjects gave informed, written consent prior to collection.[31,32] Samples were flash-frozen and stored at -80ºC until DNA extraction. Tumor stage, histology, and Barrett’s Esophagus status were determined from the pathology report. All work areas were cleaned with 70% ethanol and 10% bleach prior to DNA extraction. DNA extraction was carried out by lysing the microbes in fresh frozen tissue samples using Yeast Cell Lysis Buffer (Epicentre, Madison, WI, USA) and bead beating. Proteinase K and RNAse A were added to the samples to remove proteins and RNA, and to enrich for DNA. Samples were processed through gDNA column (Invitrogen, Carlsbad, CA, USA) and eluted in certified DNA- and RNA-free water (Mo Bio Laboratories, inc, Carlsbad, CA, USA).[25]

### University of Maryland (UMD) Esophageal Samples: PCR amplification and MiSeq sequencing

PCR amplification of the V3-4 region of the 16S rRNA gene in each sample was completed at three different dilutions of genomic DNA (1x, 10x and 100x), and the PCR reaction with the highest yield was carried forth to sequencing as previously described.[27] This process is designed to overcome the inhibitory effect of a large amount of human DNA in esophageal tissue samples. Paired-end DNA sequencing of the amplicons from all samples and both variables regions was completed in the same run on a MiSeq machine (M04141, Illumina, San Diego, CA, USA) using the 2x300 base pair chemistry (Reagent barcode: MS3917443-600V3) and 50 unique sample barcodes. All dilutions, PCR amplification and sequencing were completed at the University of Minnesota Genomics Center (Minneapolis, MN, USA).

### 16S rRNA Sequence analysis

Raw read pairs from the MiSeq platform were trimmed for quality using Trimmomatic [27] with a target final error rate of 0.5%, and merged into consensus fragments with FLASH [28]. High-quality unmerged forward reads (≥200bp after trimming) were also included for downstream analysis to increase sample coverage. PhiX spike-in fragments were detected using BLASTN [29] and removed. Sequences associated with PCR chimeras were identified using UCLUST [30] and filtered. Human genome contaminant identification was performed by aligning sequences against hg19 using Bowtie2 [31], and mitochondria and chloroplast removal utilized assignments by the RDP classifier [32]. Passing 16S rRNA gene sequences were assigned a high-resolution taxonomic lineage using Resphera Insight [33, 34].

To filter out contaminant organisms associated with DNA extraction kit reagents and other sources, we first reviewed negative controls / blank samples prepared with original tissue samples, and developed a set of dominant *indicator contaminant* species including *Bradyrhizobium spp*., *Propionibacterium_acnes, Agrobacterium_tumefaciens, Delftia spp. and Ralstonia spp*. We then performed a correlation analysis between all species/OTUs and these indicator species. Any species/OTU with a nonparametric Spearman correlation >= 0.25 was then considered to be a contaminant and was removed; however 10 species/OTUs with known human body site associations were retained including: *Faecalibacterium prausnitzii, Prevotella copri, Collinsella aerofaciens, Lactobacillus rhamnosus, Prevotella nigrescens, Prevotella disiens, and Finegoldia magna*. To filter very low frequency contaminants, we further removed all members associated with a set of genera known to be contaminants from prior literature [35]: *Bradyrhizobium, Ralstonia, Delftia, Agrobacterium, Janthinobacterium, Halomonas, Methylobacterium, Aquamicrobium, Diaphorobacter, Herbaspirillum, and Variovorax*. After contaminant removal, samples were normalized through rarefaction to 500 sequences per sample. Alpha and beta-diversity analysis performed with QIIME [36].

### Processing of The Cancer Genome Atlas (TCGA) Samples

RNA-seq and WGS bam files reflecting cancer and non-cancer samples from esophageal carcinoma patients available from TCGA were identified using the Genomic Data Commons (GDC) portal and downloaded using the GDC data transfer client (http://portal.gdc.cancer.gov/). Barrett’s esophagus status, tumor stage, gender, race and survival information were also retrieved when available from the GDC.

### Quality control and identification of microbial DNA

Unmapped sequences from the raw RNA-seq and WGS bam files were converted to FASTQ format using Samtools [37] and trimmed for quality with Trimmomatic [27] to remove error-prone reads. Additionally, in order to remove unmapped spliced transcripts and other poorly aligning sequences, we performed a local alignment to the human reference (hg19) using Bowtie2 [31]. Clean sequences passing all filters were assigned to a taxonomic lineage using Pathoscope (v1.0) [38, 39].

To filter out contaminant organisms associated with DNA extraction kit reagents and other laboratory sources, we developed a set of 10 dominant indicator contaminant species including members of Bradyrhizobium, Propionibacterium, Pseudomonas, and Arthrobacter. We then performed analysis between all species/OTUs and these indicator species across WGS tumor, WGS normal and RNA-seq samples. Any species detected in at least 58 of RNA-seq samples, or 55 of WGS tumor or 55 of WGS normal samples was often found to show a strong Spearman correlation with one or more of the indicator contaminant species, and were thus assigned putative contaminant status. We further removed all species associated with a set of higher taxa known to be contaminants from published literature [35] or that were also highly recurrent across most samples including members of Pseudomonadales, Comamonadaceae, Rhizobiales, Burkholderiales, Paenibacillaceae, Propionibacterium acnes, Escherichia, and Bacillaceae.

### Tumor Mutational Burden and Mutational Signature Analysis

We retrieved high quality nonsynonymous mutation calls from the esophageal carcinoma samples provided by the TCGA MC3 Project [40] (requiring a minimum MAF of 10% and 4 variant supporting reads) in order to calculate tumor mutational load per sample. We further extracted the trinucleotide DNA context for each SNV mutation reported to determine the underlying mutational signatures for each tumor using the *deconstructSigs* R package [41].

### Integration of 16S rRNA and TCGA Microbial Profiles

In order to provide a direct comparison between the 16S rRNA and TCGA WGS/RNA-seq microbial profiles, we first performed a concordance study at the species level across all technologies. Manual examination of the WGS/RNA-seq and 16S rRNA data revealed that some species in WGS previously determined to be contaminant were more likely to reflect true oral and upper respiratory tract species (such as *Rothia mucilaginosa* and *Streptococcus mitis*).

Therefore, we revisited the contaminant removal process for our data integration of 16S rRNA WGS / RNAseq data, and rescued species that were present in the 16S rRNA contaminant-free dataset, or those reported in a second esophageal tissue 16S rRNA study by Gall et al [42]. This effort confirmed consistent taxonomic profiles for joint interpretation across genomic data types.

### Inferred microbial metabolism

The input files were a FASTA file of representative sequences and a BIOM table of the abundance of each ASV across each sample from the NCI-MD cohort. The steps of the pipeline used were (1) sequence placement, (2) hidden-state prediction of genomes, (3) metagenome prediction, and (4) pathway-level predictions. The following pipeline was followed to perform this analysis: https://github.com/picrust/picrust2/wiki/Full-pipeline-script

### Statistical Methods

Statistical comparisons were performed in R (cran.r-project.org). Kaplan Meier analyses were performed using the *survival* and *survminer* R packages. To establish associations of specific microbial members with tumor status, we utilized Generalized linear fixed effects models (GLMs) and Generalized linear mixed effects models (GLMMs) in which patient membership was considered a fixed effort, or random effect, respectively. The Mann-Whitney test for differential abundance was applied per each genomic data type independent as a supplement to GLM analyses. Fisher’s exact test was applied to evaluate differential frequencies of positive vs negative status for each microbial member in the integrated analysis. *Generalized linear models* – taxon % abundance modeled by Tumor / Normal status (fixed effect) and Patient ID (fixed effect) (stratified by genomic data type). *Generalized linear mixed effects models* – taxon % abundance modeled by Tumor / Normal status (fixed effect) and Patient ID (random effect) (stratified by genomic data type). Fisher’s exact test for positive status (stratified by genomic data type). Comparisons to adjust for the blood-derived normal samples in TCGA were also applied. All analysis for the main figures are located at https://github.com/GreathouseLab/esoph-micro-cancer-workflow

## Data Availability

All de-identified data and code used to conduct analyses and generate figures for this manuscript are available from TCGA or at https://github.com/GreathouseLab/esoph-micro-cancer-workflow. All sequences generated during this study will be deposited on the sequence read archive prior to publication. Any protocols will be made available at the request of the researcher.

## Acknowledgements

We would like to thank all of the participants of the NCI-MD cohort study who kindly agreed to provide their data and samples for this research study.

## Funding

A. Choudhury is funded by a Postdoctoral Fellowship Award from Baylor University.

## Competing Interests

The Authors declare no Competing Financial Interests or Non-Financial Interests.

## Author Contributions

C.H and A.V. conceptualized study ideas; K.L.G, C.H., J.S. and A.V. and J.W. contributed to methodology, study design, and data curation, A.C., K.L.G., A.J., N.P., J.W. and J.S. contributed to formal analysis, bioinformatics, sequencing, visualization and validation of datasets; C.H., K.L.G., J.S., and A.V. contributed to original draft preparation and review/editing.

## Supplemental Figure Legends

**Fig. S1: Alpha and beta diversity are variable across cohorts**.

Alpha diversity within the NCI-MD and TCGA cohorts. **A-C** Violin or box plots indicate median values with upper and lower quartiles. Significance determined by Kruskal-Wallis test. **C** Alpha diversity in **B** TCGA RNA-seq and **C** WGS cohorts. Boxplots indicate median values with upper and lower quartiles. Significance determined by Wilcoxon test. **D-E** Beta diversity within the NCI-MD case control study determined by Bray-Curtis. **E** Compositional PCoA biplot of beta diversity with arrows signifying highly ranked taxonomic features contributing the most difference. Significance determined by PERMANOVA test. For all graphs, n.s. not significant, * *p* < 0.05.

**Fig. S2: Taxa abundance are not associated with risk factors for esophageal cancer in NCIMD cohort**. Taxa abundance in NCI-MD case control study for **A** tissue type, **B** gender, **C** histology, **D** race, **E** smoking status, and **F** stage. For all graphs, violin plots indicate median values with upper and lower quartiles. Significance was determined by Kruskal-Wallis test n.s. not significant, * *p* < 0.05, ** *p* < 0.01, *** *p* < 0.001.

**Figure S3 - Taxa abundance are not associated with *TP53* mutation status in ESCA TCGA WGS and RNA-seq cohort**. Taxa abundance TCGA WGS study for TP53 mutation status in **A** WGS and **B** RNA-seq dataset. For all graphs, violin plots indicate median values with upper and lower quartiles. Significance was determined by Kruskal-Wallis test n.s. not significant, * *p* < 0.05, ** *p* < 0.01, *** *p* < 0.001.

**Figure. S4: Heatmap of taxa co-enrichment networks as determined by SparCC**.

**A** Taxa enrichment heatmap for NCI-MD case control study with underlying data for network depicted in Fig. 3A. **B** Taxa enrichment heatmap for TCGA RNA-seq with underlying data for network depicted in Fig. 3B. **C** Taxa enrichment heatmap for TCGA WGS with underlying data for network depicted in Fig. 3C. Data was calculated for each heatmap as described in Fig. 3.

**Figure. S5: Heatmap of immune cells significantly enriched or depleted in tumors with high carriage of ESCA-enriched taxa**.

**A** RNA-sequencing was performed on NCI-MD patients (n = 27; non-tumor = 13, BO = 4, tumor = 10) and samples were analyzed for predicted cell infiltration using xCell (citation). Cell infiltrates and taxa abundance were correlated using Spearman’s coefficient. **B** Correlation of xCell predicted cell infiltration in TCGA RNA-seq patients with taxa abundance. All correlations shown were sig. *p* < 0.05.

**Figure. S6: MEP infiltration is lower in taxa-positive tumors**.

Quantification of predicted megakaryocyte-erythroid progenitor (MEP) cell infiltration in the tumor tissues of NCI-MD case control study (top) and TCGA RNA-seq (bottom) present or absent for the indicated taxa. Tumors classified as “present” had one or more reads for the given taxa. MEP infiltration predicted by xCell as described in Fig. 4. Significance determined by Wilcoxon test, n.s. not significant, * *p* < 0.05, ** *p* < 0.01, *** *p* < 0.001.

**Figure. S7: Platelet infiltration trends higher in taxa-positive tumors**.

Quantification of predicted platelet infiltration in the tumor tissues of NCI-MD case control study (top) and TCGA RNA-seq (bottom) present or absent for the indicated taxa. Tumors classified as “present” had one or more reads for the given taxa. Platelet infiltration predicted by xCell as described in Fig. 4. Significance determined by Wilcoxon test, n.s. not significant, * *p* < 0.05.

**Table S1-S3. Patient demographics by cohort overview**.

**Table S4. Concordance between taxonomic abundance across cohorts**.

**Table S5. Concordance between taxonomic co-enrichment and co-exclusion across cohorts**.

## Notes

https://github.com/GreathouseLab/esoph-micro-cancer-workflow

### Competing Interest Statement

The authors have declared no competing interest.

