## Supplemental Figures and Tables for "Co-enrichment of cancer-associated bacterial taxa is correlated with immune cell infiltrates in esophageal tumor tissue": Supplemental_Figure_FINAL.pdf

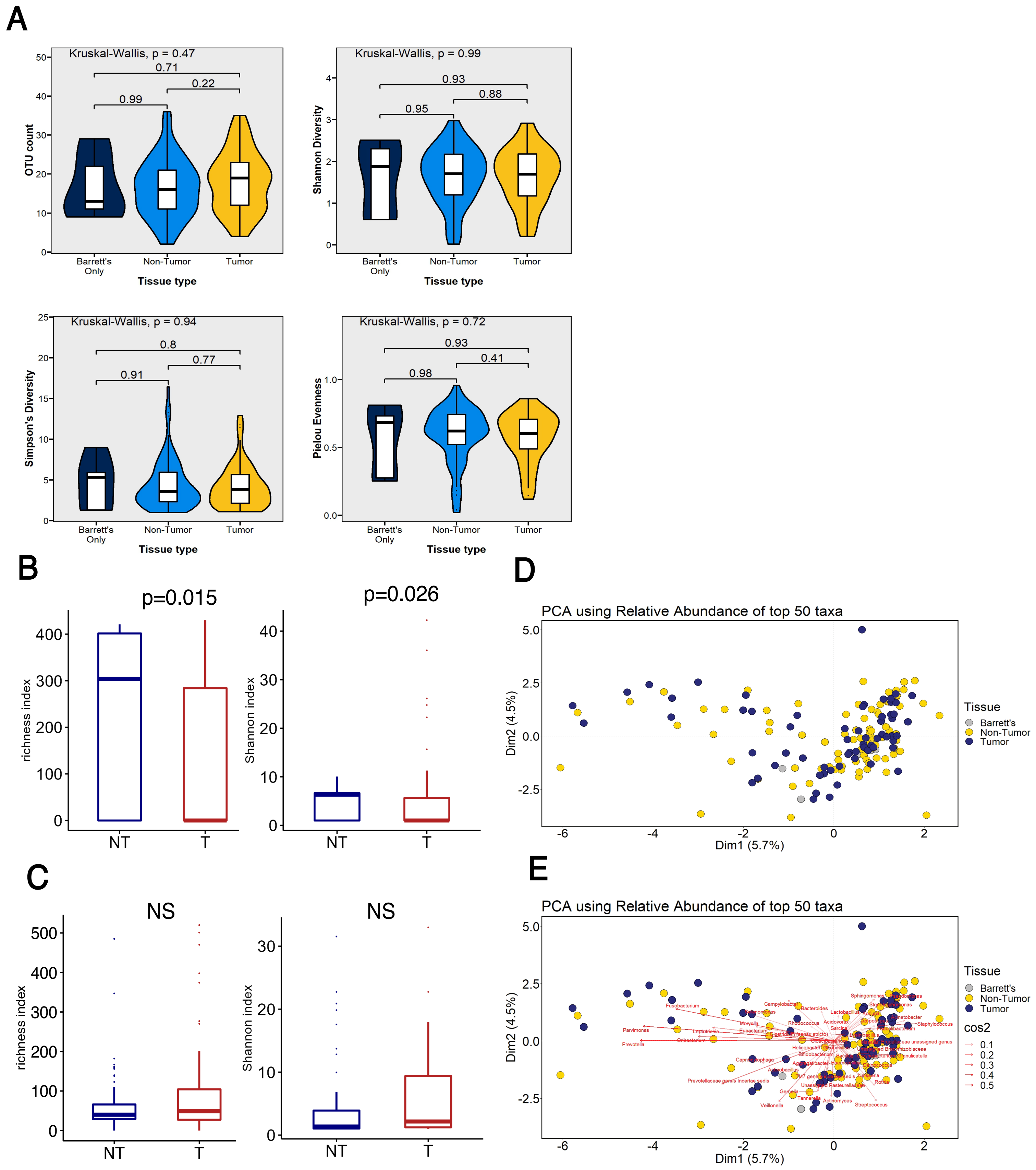

A
Kruskal-Wallis, p = 0.47
0.71
0.22
0.99
OTU count
Kruskal-Wallis, p = 0.99
0.93
0.88
0.95
Shannon Diversity
Kruskal-Wallis, p = 0.94
0.8
0.77
0.91
Simpson's Diversity
Kruskal-Wallis, p = 0.72
0.93
0.41
0.98
Pielou Evenness
Barrett's Only
Non-Tumor
Tumor
Tissue type
B
p=0.015
richness index
p=0.026
Shannon index
NT
T
C
NS
richness index
NS
Shannon index
NT
T
D
PCA using Relative Abundance of top 50 taxa
Dim1 (5.7%)
Dim2 (4.5%)
Tissue
Barrett's
Non-Tumor
Tumor
E
PCA using Relative Abundance of top 50 taxa
Dim1 (5.7%)
Dim2 (4.5%)
Tissue
Barrett's
Non-Tumor
Tumor
cos2
0.1
0.2
0.3
0.4
0.5


Figure S1

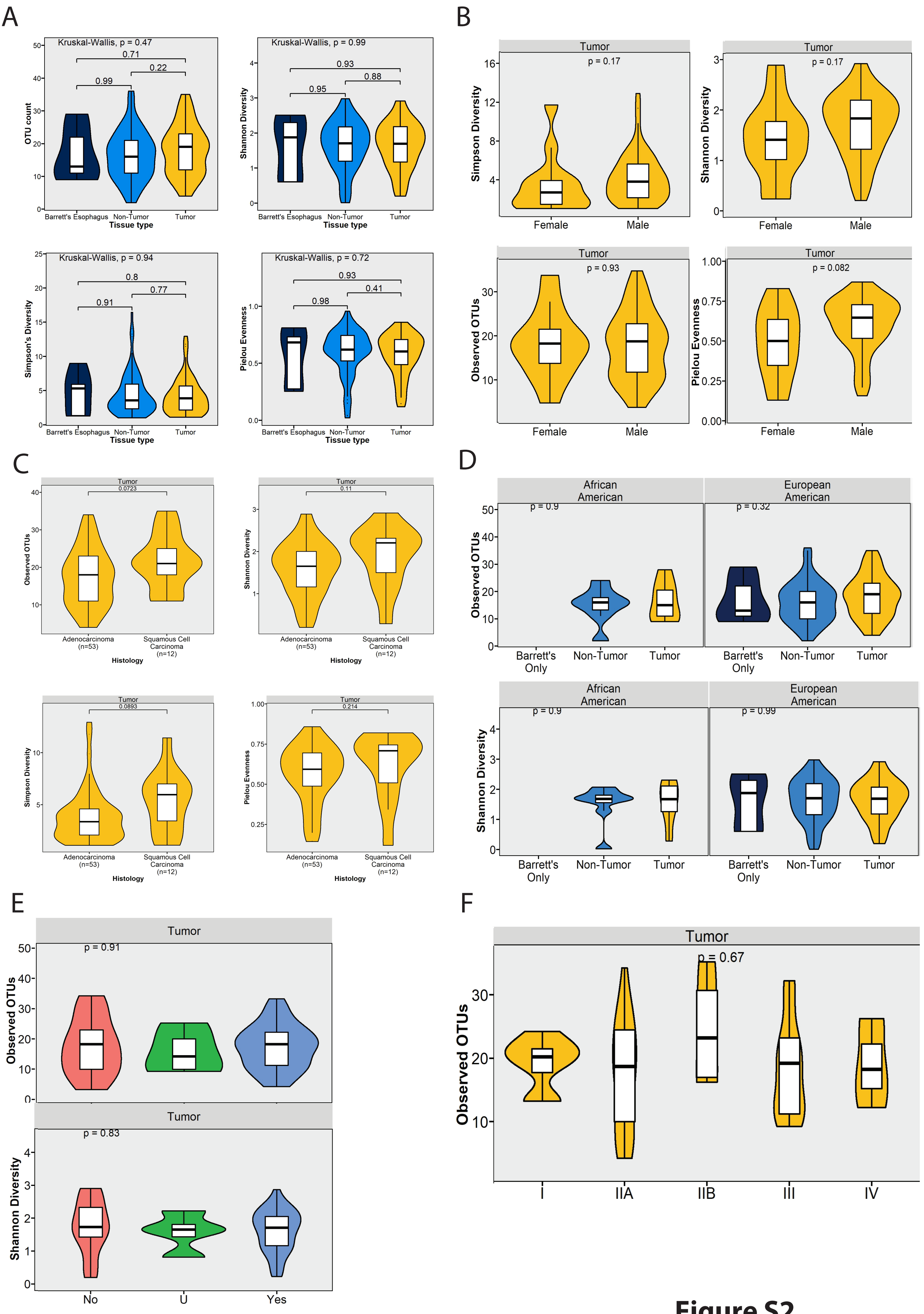
A
B
C
D
E
F
Kruskal-Wallis, p = 0.47
OTU count
Tissue type
Barrett's Esophagus
Non-Tumor
Tumor
Kruskal-Wallis, p = 0.99
Shannon Diversity
Kruskal-Wallis, p = 0.94
Simpson's Diversity
Kruskal-Wallis, p = 0.72
Pielou Evenness
Simpson Diversity
Shannon Diversity
Observed OTUs
Pielou Evenness
Female
Male
Histology
Adenocarcinoma (n=53)
Squamous Cell Carcinoma (n=12)
African American
European American
Barrett's Only
No
U
Yes
I
IIA
IIB
III
IV

**Figure S2**

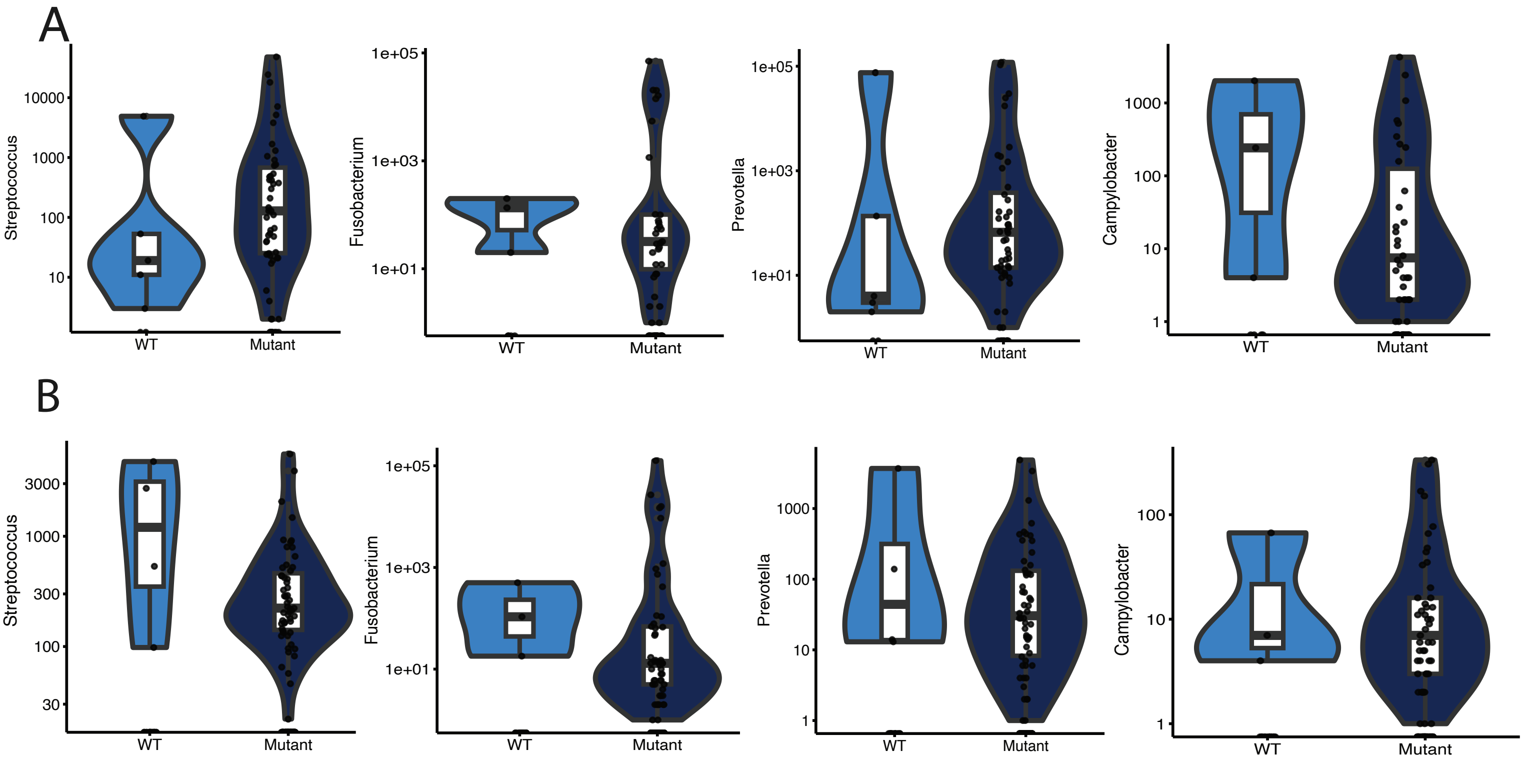
A
Streptococcus
10000
1000
100
10
WT
Mutant
Fusobacterium
1e+05
1e+03
1e+01
WT
Mutant
Prevotella
1e+05
1e+03
1e+01
WT
Mutant
Campylobacter
1000
100
10
1
WT
Mutant
B
Streptococcus
3000
1000
300
100
30
WT
Mutant
Fusobacterium
1e+05
1e+03
1e+01
WT
Mutant
Prevotella
1000
100
10
1
WT
Mutant
Campylobacter
100
10
1
WT
Mutant

Figure S3.

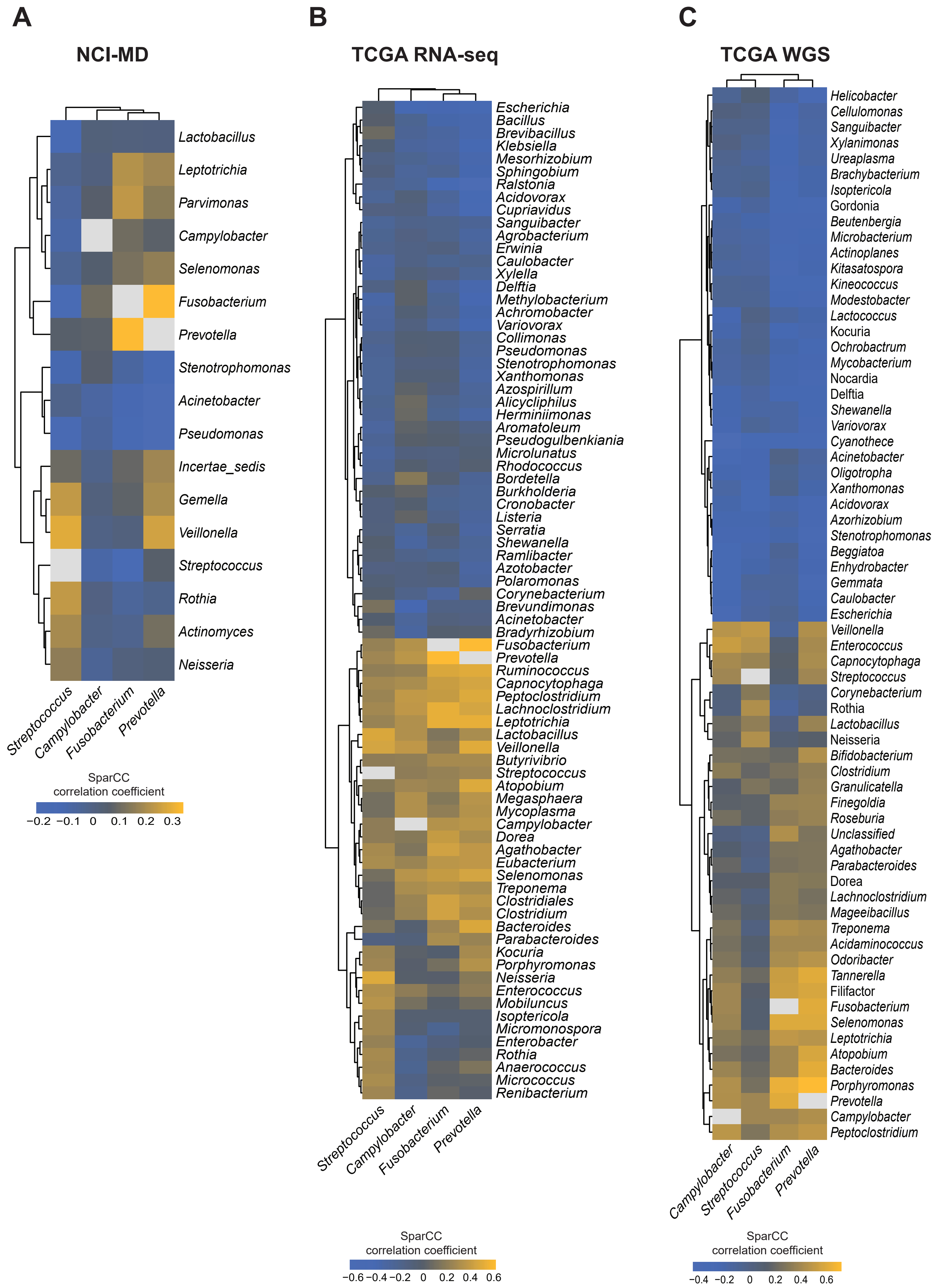
A
NCI-MD
Lactobacillus
Leptotrichia
Parvimonas
Campylobacter
Selenomonas
Fusobacterium
Prevotella
Stenotrophomonas
Acinetobacter
Pseudomonas
Incertae_sedis
Gemella
Veillonella
Streptococcus
Rothia
Actinomyces
Neisseria
Streptococcus
Campylobacter
Fusobacterium
Prevotella
SparCC
correlation coefficient
−0.2 −0.1 0 0.1 0.2 0.3
B
TCGA RNA-seq
Escherichia
Bacillus
Brevibacillus
Klebsiella
Mesorhizobium
Sphingobium
Ralstonia
Acidovorax
Cupriavidus
Sanguibacter
Agrobacterium
Erwinia
Caulobacter
Xylella
Delftia
Methylobacterium
Achromobacter
Variovorax
Collimonas
Pseudomonas
Stenotrophomonas
Xanthomonas
Azospirillum
Alicycliphilus
Herminiimonas
Aromatoleum
Pseudogulbenkiania
Microlunatus
Rhodococcus
Bordetella
Burkholderia
Cronobacter
Listeria
Serratia
Shewanella
Ramlibacter
Azotobacter
Polaromonas
Corynebacterium
Brevundimonas
Acinetobacter
Bradyrhizobium
Fusobacterium
Prevotella
Ruminococcus
Capnocytophaga
Peptoclostridium
Lachnoclostridium
Leptotrichia
Lactobacillus
Veillonella
Butyrivibrio
Streptococcus
Atopobium
Megasphaera
Mycoplasma
Campylobacter
Dorea
Agathobacter
Eubacterium
Selenomonas
Treponema
Clostridiales
Clostridium
Bacteroides
Parabacteroides
Kocuria
Porphyromonas
Neisseria
Enterococcus
Mobiluncus
Isoptericola
Micromonospora
Enterobacter
Rothia
Anaerococcus
Micrococcus
Renibacterium
Streptococcus
Campylobacter
Fusobacterium
Prevotella
SparCC
correlation coefficient
−0.6 −0.4 −0.2 0 0.2 0.4 0.6
C
TCGA WGS
Helicobacter
Cellulomonas
Sanguibacter
Xylanimonas
Ureaplasma
Brachybacterium
Isoptericola
Gordonia
Beutenbergia
Microbacterium
Actinoplanes
Kitasatospora
Kineococcus
Modestobacter
Lactococcus
Kocuria
Ochrobactrum
Mycobacterium
Nocardia
Delftia
Shewanella
Variovorax
Cyanothece
Acinetobacter
Oligotropha
Xanthomonas
Acidovorax
Azorhizobium
Stenotrophomonas
Beggiatoa
Enhydrobacter
Gemmata
Caulobacter
Escherichia
Veillonella
Enterococcus
Capnocytophaga
Streptococcus
Corynebacterium
Rothia
Lactobacillus
Neisseria
Bifidobacterium
Clostridium
Granulicatella
Finegoldia
Roseburia
Unclassified
Agathobacter
Parabacteroides
Dorea
Lachnoclostridium
Mageeibacillus
Treponema
Acidaminococcus
Odoribacter
Tannerella
Filifactor
Fusobacterium
Selenomonas
Leptotrichia
Atopobium
Bacteroides
Porphyromonas
Prevotella
Campylobacter
Peptoclostridium
Campylobacter
Streptococcus
Fusobacterium
Prevotella
SparCC
correlation coefficient
−0.4 −0.2 0 0.2 0.4 0.6

**Figure S4**

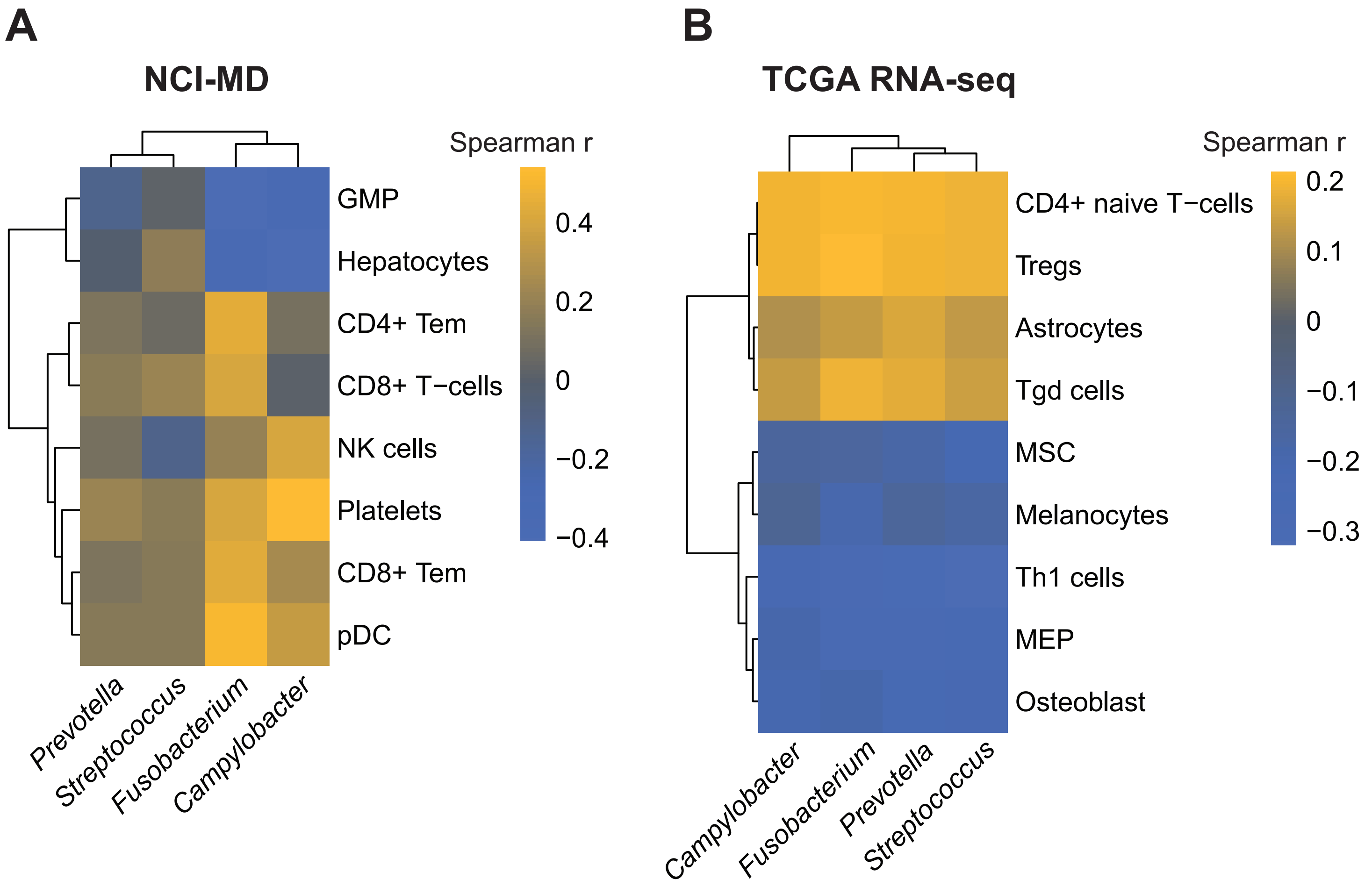
A
NCI-MD
Spearman r
0.4
0.2
0
−0.2
−0.4
GMP
Hepatocytes
CD4+ Tem
CD8+ T−cells
NK cells
Platelets
CD8+ Tem
pDC
Prevotella
Streptococcus
Fusobacterium
Campylobacter
B
TCGA RNA-seq
Spearman r
0.2
0.1
0
−0.1
−0.2
−0.3
CD4+ naive T−cells
Tregs
Astrocytes
Tgd cells
MSC
Melanocytes
Th1 cells
MEP
Osteoblast
Campylobacter
Fusobacterium
Prevotella
Streptococcus

**Figure S5**

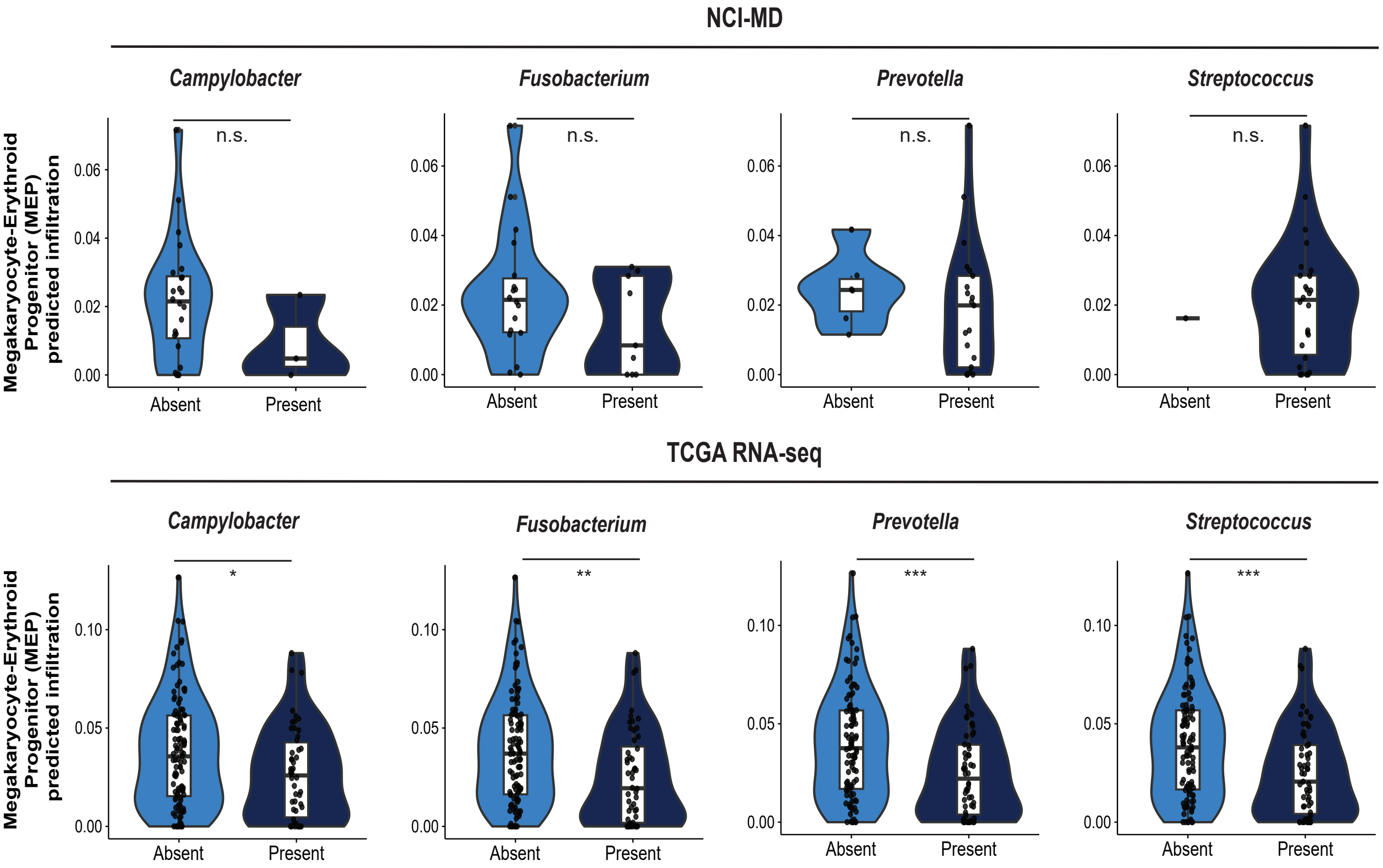

NCI-MD
Campylobacter
Fusobacterium
Prevotella
Streptococcus
n.s.
Megakaryocyte-Erythroid Progenitor (MEP) predicted infiltration
0.00
0.02
0.04
0.06
Absent
Present
TCGA RNA-seq
*
**
***
0.05
0.10


Figure S6

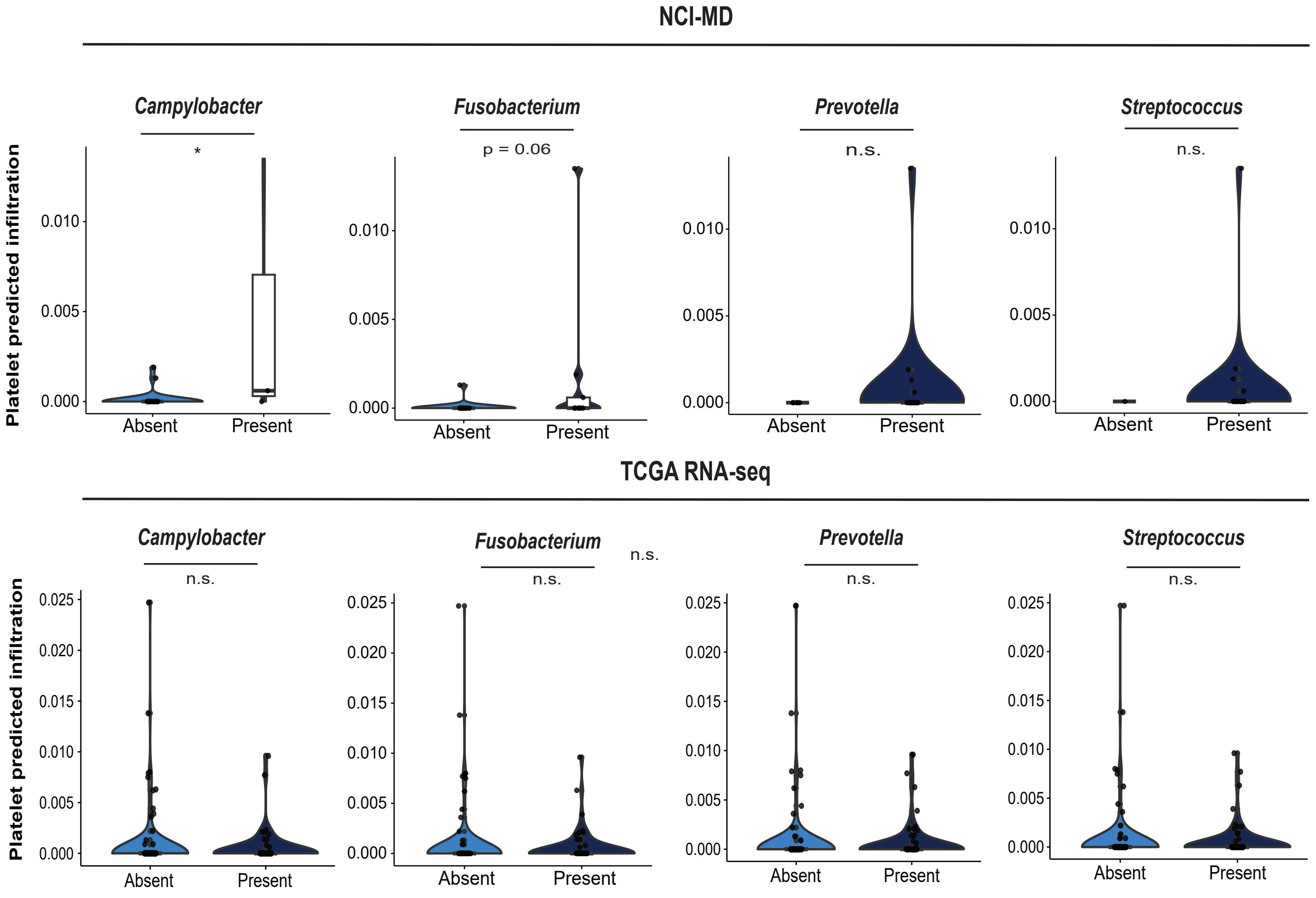
NCI-MD
Campylobacter
*
Fusobacterium
p = 0.06
Prevotella
n.s.
Streptococcus
n.s.
Platelet predicted infiltration
0.010
0.005
0.000
Absent
Present
TCGA RNA-seq
Campylobacter
n.s.
Fusobacterium
n.s.
n.s.
Prevotella
n.s.
Streptococcus
n.s.
0.025
0.020
0.015
0.010
0.005
0.000
Absent
Present

Figure S7

| Primary Cohort (NCI-MD Samples) | | | | | | |
|---|---|---|---|---|---|---|
| **Description** | **Esophageal adeno-carcinoma tissues** | **Esophageal squamous cell carcinoma tissue** | **Esophageal adeno-carcinoma-adjacent tissues** | **Esophageal squamous cell carcinoma-adjacent tissue** | **Barrett's Esophagus tissue** | **Total** |
| **N samples** | 74 | 17 | 87 | 27 | 8 | 213 |
| **Sex (N, %)** | | | | | | |
| Male | 65 (87.8) | 9 (52.9) | 77 (88.5) | 14 (51.9) | 8 (100) | 173 (81.2) |
| Female | 9 (12.2) | 8 (47.1) | 10 (11.5) | 13 (48.1) | 0 (0.0) | 40 (18.8) |
| **Race (N, %)** | | | | | | |
| European American | 72 (97.3) | 10 (58.8) | 85 (97.7) | 16 (59.3) | 8 (100) | 191 (89.7) |
| African American | 1 (1.4) | 6 (35.3) | 1 (1.1) | 9 (33.3) | 0 (0.0) | 17 (8.0) |
| Other | 1 (1.4) | 1 (5.9) | 1 (1.1) | 2 (7.4) | 0 (0.0) | 5 (2.3) |
| **Age, mean ± SD** | 60.3 +/- 10.6 | 60.7 +/- 9.3 | 61.9 +/- 11.0 | 58.7 +/- 9.2 | 65.9 +/- 6.0 | 61.0 +/- 10.4 |
| **BMI (N, %)** | | | | | | |
| underweight | 1 (1.4) | 2 (11.8) | 1 (1.1) | 5 (18.5) | 0 (0.0) | 9 (4.2) |
| normal weight | 21 (28.4) | 9 (52.9) | 19 (21.8) | 11 (40.7) | 1 (12.5) | 61 (28.6) |
| overweight | 24 (32.4) | 5 (29.4) | 27 (31.0) | 6 (22.2) | 2 (25.0) | 64 (30.0) |
| obese | 23 (31.1) | 1 (5.9) | 27 (31.0) | 2 (7.4) | 3 (37.5) | 56 (26.3) |
| unknown | 5 (6.8) | 0 (0.0) | 13 (14.9) | 3 (11.1) | 2 (25.0) | 23 (10.8) |
| **History of Barrett's Esophagus (N, %)** | | | | | | |
| yes | 50 (67.6) | 3 (17.6) | 60 (69.0) | 4 (14.8) | 8 (100) | 125 (58.7) |
| no | 24 (32.4) | 14 (82.4) | 27 (31.0) | 23 (85.2) | 0 (0.0) | 88 (41.3) |
| **History of smoking (N,%)** | | | | | | |
| yes | 56 (75.7) | 14 (82.4) | 64 (73.6) | 22 (81.5) | 5 (62.5) | 161 (75.6) |
| no | 15 (20.3) | 2 (11.8) | 14 (16.1) | 2 (7.4) | 2 (25.0) | 35 (16.4) |
| missing | 3 (4.1) | 1 (5.9) | 9 (10.3) | 3 (11.1) | 1 (12.5) | 17 (8.0) |
| **Clinical Stage (N, %)** | | | | | | |
| 0 | 2 (2.7) | 0 (0.0) | 3 (3.4) | 0 (0.0) | | |
| I | 35 (47.3) | 1 (5.9) | 12 (13.8) | 1 (3.7) | | |

| | | | | |
|---|---|---|---|---|
| II | 5 (6.8) | 6 (35.3) | 36 (41.4) | 12 (44.4) |
| III | 26 (35.1) | 1 (5.9) | 28 (32.2) | 12 (44.4) |
| IV | 5 (6.8) | 9 (52.9) | 7 (8.0) | 2 (7.4) |
| missing | 0 (0.0) | 0 (0.0) | 0 (0.0) | 0 (0.0) |
| **Neoadjuvant therapy (N, %)** | 44 (59.5) | 9 (52.9) | 52 (59.8) | 18 (66.7) |
| **Survival (days), mean ± SD** | 1922.9 +/- 1813.0 | 1924.0 +/- 1872.0 | 1848.7 +/- 1754.4 | 1699 .2+/- 1911.4 |

### Table S1. NCIMD case control cohort study sample demographics

| | Mean % Abundance in Tumor Samples | | |
|---|---|---|---|
| Species | 16S rRNA | WGS | RNAseq |
| *Streptococcus spp.** | 20.17 | 15.52 | 1.11 |
| *Prevotella melaninogenica* | 1.99 | 10.25 | 0.34 |
| *Fusobacterium nucleatum* | 4.38 | 6.31 | 2.07 |
| *Veillonella parvula* | 1.34 | 4.61 | 0.42 |
| *Helicobacter pylori* | 0.18 | 3.23 | 0.01 |
| *Haemophilus influenzae* | 0.22 | 2.86 | 0.66 |
| *Prevotella denticola* | 0.13 | 2.22 | 0.17 |
| *Prevotella intermedia* | 0.80 | 1.81 | 0.05 |
| *Campylobacter concisus* | 0.14 | 1.60 | 0.03 |
| *Staphylococcus aureus* | 0.54 | 1.23 | 0.16 |
| *Rothia mucilaginosa* | 3.16 | 1.09 | 0.54 |
| *Haemophilus parainfluenzae* | 1.74 | 1.09 | 0.09 |
| *Selenomonas sputigena* | 1.96 | 1.01 | 0.08 |
| *Bacteroides fragilis* | 0.16 | 0.77 | 0.03 |
| *Stenotrophomonas maltophilia* | 3.88 | 0.47 | 0.41 |
| *Gemella haemolysans* | 1.55 | 0.32 | 0.16 |

*S. oralis, S. mitis, S. pneumoniae, S. parasanguinis, S. salivarius*

| | | Comparative Statistical Results | | | | | Statistical Frequency Results | | |
|---|---|---|---|---|---|---|---|---|---|
| | | GLM P-value | GLMM P-value | Mann-Whitney Tumor vs Non-Tumor | | | Fisher's Exact Test for Tumor vs Normal | | |
| Species | Cohort Exclusion | Tumor vs Non-tumor | Tumor vs Non-tumor | 16SrRNA | WGS | RNAseq | 16SrRNA | WGS | RNAseq |
| *Fusobacterium nucleatum* | No WGS Blood | 0.0870 | 0.0054 | 0.0000 | 0.4602 | 0.0524 | 0.0004 | 0.8134 | 0.0471 |
| *Fusobacterium nucleatum* | WGS Non-tumor w/ Blood | 0.0015 | 0.0001 | 0.0000 | 0.0074 | 0.0524 | 0.0004 | 0.0253 | 0.0471 |

| | | Comparative Statistical Results | | | | | Statistical Frequency Results | | |
|---|---|---|---|---|---|---|---|---|---|
| | | GLM P-value | GLMM P-value | Mann-Whitney Tumor vs Non-Tumor | | | Fisher's Exact Test for Tumor vs Normal | | |
| Species | Cohort Exclusion | Tumor vs Non-tumor | Tumor vs Non-tumor | 16SrRNA | WGS | RNAseq | 16SrRNA | WGS | RNAseq |
| *Streptococcus spp* | No WGS Blood | 0.3952 | 0.5795 | 0.0154 | 0.6974 | 0.2194 | 0.0228 | 0.7872 | 0.1066 |
| *Streptococcus spp* | WGS Non-tumor w/ Blood | 0.0873 | 0.0151 | 0.0154 | 0.0346 | 0.2194 | 0.0228 | 0.0127 | 0.1066 |

| | | Comparative Statistical Results | | | | | Statistical Frequency Results | | |
|---|---|---|---|---|---|---|---|---|---|
| | | GLM P-value | GLMM P-value | Mann-Whitney Tumor vs Non-Tumor | | | Fisher's Exact Test for Tumor vs Normal | | |
| Species | Cohort Exclusion | Tumor vs Non-tumor | Tumor vs Non-tumor | 16SrRNA | WGS | RNAseq | 16SrRNA | WGS | RNAseq |
| *Campylobacter concisus* | No WGS Blood | 0.2805 | 0.4084 | 0.0047 | 0.6204 | 0.0388 | 0.0045 | 1.0000 | 0.0419 |
| *Campylobacter concisus* | WGS Non-tumor w/ Blood | 0.2148 | 0.1485 | 0.0047 | 0.0077 | 0.0388 | 0.0045 | 0.0089 | 0.0419 |

| | | Comparative Statistical Results | | | | | Statistical Frequency Results | | |
|---|---|---|---|---|---|---|---|---|---|
| | | GLM P-value | GLMM P-value | Mann-Whitney Tumor vs Non-Tumor | | | Fisher's Exact Test for Tumor vs Normal | | |
| Species | Cohort Exclusion | Tumor vs Non-tumor | Tumor vs Non-tumor | 16SrRNA | WGS | RNAseq | 16SrRNA | WGS | RNAseq |
| *Prevotella melaninogenica* | No WGS Blood | 0.9021 | 0.8310 | 0.9459 | 0.1724 | 0.0797 | 1.0000 | 0.1254 | 0.1884 |
| *Prevotella melaninogenica* | WGS Non-tumor w/ Blood | 0.0002 | 0.0006 | 0.9459 | 0.0000 | 0.0797 | 1.0000 | 0.0000 | 0.1884 |

**Table S4.** Mean relative abundance across cohorts sorted first by species in WGS and comparative statical comparisons of four taxa enriched in tumors vs non-tumor adjacent.

| | *Campylobacter* | | | *Fusobacterium* | | | *Prevotella* | | | *Streptococcus* | | |
|---|---|---|---|---|---|---|---|---|---|---|---|---|
| **Genus** | **NCIMD** | **RNA** | **WGS** | **NCIMD** | **RNA** | **WGS** | **NCIMD** | **RNA** | **WGS** | **NCIMD** | **RNA** | **WGS** |
| *Campylobacter* | NA | NA | NA | 0.0977 | 0.3314 | 0.3733 | 0.0692 | 0.3494 | 0.4239 | -0.0923 | 0.1915 | 0.3829 |
| *Fusobacterium* | 0.0977 | 0.3314 | 0.3733 | NA | NA | NA | 0.3371 | 0.609 | 0.6038 | -0.2083 | 0.2132 | 0.0937 |
| *Leptotrichia* | 0.0072 | 0.3211 | 0.3086 | 0.2125 | 0.5202 | 0.4781 | 0.1792 | 0.5202 | 0.4643 | -0.0326 | 0.2232 | 0.2 |
| *Neisseria* | -0.0485 | -0.0019 | | 0.0138 | -0.0152 | | 0.0233 | 0.1697 | | 0.1532 | 0.4782 | |
| *Prevotella* | 0.0692 | 0.3494 | 0.4239 | 0.3371 | 0.609 | 0.6038 | NA | NA | NA | 0.0637 | 0.2522 | 0.3786 |
| *Selenomonas* | 0.0531 | | 0.3642 | 0.1159 | | 0.5826 | 0.1559 | | 0.5889 | -0.0460 | | 0.0997 |
| *Streptococcus* | -0.0923 | 0.1915 | 0.3829 | -0.2083 | 0.2132 | 0.0937 | 0.0637 | 0.2522 | 0.3786 | NA | NA | NA |
| *Veillonella* | -0.0134 | 0.3612 | 0.4588 | 0.0137 | 0.1892 | -0.123 | 0.2569 | 0.4912 | 0.4286 | 0.2815 | 0.4452 | 0.4925 |

**Table S5.** Concordance between taxonomic co-occurrence and co-exclusion
